# Acute deletion of PLIN5 in brown adipocytes causes mitochondrial dysfunction and cold intolerance

**DOI:** 10.64898/2025.12.31.697198

**Authors:** Violeta I. Gallardo Montejano, Chaofeng Yang, Hannah Hurtado, Perry E. Bickel

## Abstract

Cold exposure of mice is associated with adaptive molecular and organellar changes in brown adipose tissue (BAT) that promote thermogenesis to defend body temperature. We previously reported that the lipid droplet protein Perilipin 5 (PLIN5) robustly increases in BAT during acute exposure of mice to cold. We demonstrated that chronic induction of BAT PLIN5 within the physiological range in mice housed at room temperature mimics the effects of cold exposure in terms of increased thermogenic gene expression in BAT, increased BAT mitochondria cristae packing, and increased uncoupled mitochondrial respiration. Additionally, BAT PLIN5 overexpression led to healthy remodeling of inguinal white adipose tissue with improved systemic glucose tolerance and reduced diet-induced hepatic steatosis. Conversely, PLIN5 constitutive deletion in brown adipose tissue resulted in decreased BAT thermogenic gene expression and in BAT mitochondrial dysfunction but did not lead to cold intolerance or changes in glucose tolerance. We hypothesized that preserved cold tolerance despite chronic deficiency of PLIN5 in BAT was the result of compensatory white adipose tissue (WAT) beiging, as suggested by the observed increase in thermogenic gene expression in inguinal WAT (iWAT). To test this hypothesis, we developed a mouse model of doxycycline-inducible, acute deficiency of PLIN5 in BAT of adult mice (BiKOPLIN5 mice). After 7 days of doxycycline treatment and housing at 6 °C, PLIN5 was significantly reduced in the BAT of BiKOPLIN5 mice compared with littermate control mice but was unchanged in the iWAT of these experimental groups. Under these conditions, thermogenic gene expression was reduced significantly in the BAT of BiKOPLIN5 mice compared to Control mice, as were mitochondrial cristae density and uncoupled BAT mitochondrial respiration. These effects of acute PLIN5 deficiency in BAT were associated with cold intolerance, which was consistent with the observed failure in iWAT of thermogenic gene expression to increase beyond that of Controls. These findings clarify the essential role of BAT PLIN5 in the physiological adaptive responses of mice to cold ambient temperature.

## Introduction

PLIN5 is a lipid droplet protein that is expressed in highly oxidative tissues such as heart, oxidative skeletal muscle, fasted liver, and brown adipose tissue (BAT) [1–3]. Under basal conditions, PLIN5 resides on the surface of lipid droplets and in the cytosol. Upon activation of the β-adrenergic-protein kinase A (PKA) pathway by catecholamines, Mouse PLIN5 is phosphorylated on serine 155 and exerts distinct functions on the lipid droplet and in the nucleus. On the lipid droplet surface PLIN5 coordinates the activation of adipose triglyceride lipase (ATGL) to hydrolyze the triglycerides sequestered within droplets, [4]. Once phosphorylated PLIN5 also enriches in the nucleus where it promotes the SIRT1-PGC1α gene program to augment mitochondrial biogenesis and function [5]. Additionally, PLIN5 is proposed to associate with mitochondria to form a physical tether between lipid droplets and mitochondria [6] [7].

Over the past few years, we have investigated the role of PLIN5 in BAT, based on its potential to promote mitochondrial oxidative capacity in that tissue. First, to explore the normal physiological role of BAT PLIN5 in the context adaptive thermogenesis, we housed C57BL/6 mice at 6 °C for intervals of up to 2 weeks. We reported that PLIN5 protein and mRNA increased significantly in BAT during cold housing and reached a maximum at 48 h [8]. Conversely, PLIN5 mRNA and protein levels were suppressed in mouse BAT after housing at thermoneutrality (30 °C) for 48 h. To dissect the function of increasing PLIN5 expression in BAT independent of cold challenge, we created a mouse model with doxycycline-inducible expression of a *Plin5* transgene. We found that a 4- to 5-fold increase in PLIN5 protein in mice housed at room temperature was associated with increased mitochondria cristae density, mitochondrial DNA, and mitochondrial respiratory capacity, as well as other markers of mitochondrial biogenesis and function. These changes in BAT mitochondria coincided with improved acute cold tolerance and chronic cold acclimation, increased systemic glucose tolerance and insulin sensitivity, decreased high fat diet-induced hepatic steatosis, and healthy inguinal white adipose tissue (iWAT) remodeling [8]. As might be predicted from these gain-of-function experiments, we reported that constitutive knockout *Plin5* in BAT via a Ucp1-Cre allele [8] resulted in dysfunctional mitochondria in BAT at room temperature and dysmorphic mitochondria with dramatically reduced cristae density in the BAT of cold-exposed mice. However, mice with constitutive knockout of *Plin5* in BAT did not exhibit cold or glucose intolerance [8] likely explained by the induction of beiging genes in iWAT with compensatory beige adipocyte thermogenesis. Herein, we report our findings in a mouse model of acute disruption of the *Plin5* gene specifically in BAT that we created to avoid the potential metabolic compensation of iWAT beiging that may have obscured loss-of-function systemic phenotypes of BAT *Plin5* constitutive gene knockout.

## Materials and Methods

### Animal studies

We performed all animal experiments with approval from the University of Texas Southwestern Medical Center (UTSWMC) Institutional Animal Care and Use Committee, and all experiments were performed in adherence to the guidelines of National Research Council, 2011, Guide for the Care and Use of Laboratory Animals: Eighth Edition [9].

For all experiments presented in this study, we used male or female mice as indicated on a C57BL/6J background. We housed mice in a conventional animal facility at 23 °C in a 12-h light/dark cycle with free access to food and water, unless otherwise indicated in the text, figure legends, or “Methods”. For controlled temperature experiments, we housed mice in a thermoneutrality box at 30 °C (Powers Scientific Inc., Model # RIS70SD) or cold box (Powers Scientific Inc., Model # RIS70SD) at 6 °C. Mouse euthanasia was by isoflurane anesthesia followed by cervical dislocation.

### Generation of mouse lines

#### BKOPLIN5 strain

We used this line in this manuscript only in Figure 6 and the characterization of this line was reported previously by us [8]. This line is a constitutive knockout line specific for *Plin5* gene deletion in BAT. To create a conditional *Plin5* allele in mice, two loxP sites were introduced flanking exons 3–8 of the *Plin5* gene (NM_025874.3). An FRT-PKG-Neo-FRT cassette [10] followed the loxP site flanking exon 8, which generated a knockout-first allele. BAC injection of the targeting construct and homologous recombination in C57BL/6J ES cells was performed by the UTSW Transgenic Core. The correct ES cell clones were screened and verified by Southern blot. The founder was backcrossed to the C57BL/6J strain. The knockout-first allele was crossed with flp mice (JAX 009086) to remove the Neo cassette and generate the floxed line, *Plin5^loxp/loxp^*. The final knockout allele (deletion of exons 3–8) was generated by crossing *Plin5^loxp/loxp^* mice with Ucp1-Cre mice to generate the BKOPLIN5 strain. The Ucp1-Cre mouse line (B6.FVB-Tg (Ucp1-Cre)1Evdr/J) was generated by the Evan Rosen Lab [11] and obtained from Jackson Laboratories Stock # 024670.

#### BiKOPLIN5 strain

This line is a doxycycline-inducible PLIN5 knockout specific for BAT. To create the BiKOPLIN5, we used our previously described *Plin5^loxp/loxp^* mice described above. We then crossed this line with transgenic mice expressing the “tet-on” transcription factor rtTA under the control of the Ucp1 gene promoter (UCP1-rtTA), which was generously provided by Philipp Scherer [12]. Finally, we crossed this line (*Plin5 ^loxp/loxp^*; Ucp1-rtTA) with a tet-responsive Cre-recombinase (TRE-Cre) line that can be activated with rtTA in the presence of doxycycline, which was obtained from Jackson Laboratories (RRID:IMSR_JAX:006234) and described previously [13].

#### Diets and timeline for experiments

For the BKOPLIN5 mice, we used chow diet (Teklad, 2016). For BiKOPLIN5 mice, we used special diets that contained 600 mg of Dox/kg diet and are referred to as Chow (S4107, BioServ) or HFD (60% high fat, S7067, BioServ). For the BiKOPLIN5 mice experiments we performed the experiments after 7 days or 21 days on Dox diet and at the indicated housing temperature.

#### Cold housing and cold tolerance test

For cold housing we single-housed the mice in a cold box (Powers Scientific Inc., Model # RIS70SD) at 6 °C. Mice had free access to food and water except during the 8-hour cold tolerance test (see below). We housed mice for 7 days or 21 days as described in the Figures and Figure legends. For cold tolerance we followed the method previously described [8] with minor modification as follows: we measured body temperature using an implantable temperature transponder (IPTT300, BioMedic Data Systems Inc, Seaford, DE) inserted subcutaneously in the dorsal side (back) of the mice, but positioned outside from the interscapular BAT region. We used the manufacturer’s needle assembly under general anesthesia with isoflurane via precision vaporizer. To allow recovery, we performed temperature experiments 3 days after the transponder insertion. After recovery we single-housed the mice, moved them to the cold box, and started 600 mg Dox chow diet. Mice were then housed for 7 days at 6 °C with free access to food and water. On day 8 we fasted the mice at 8 a.m., and we measured body temperature using a temperature reader (DAS-8007-IUS, BioMedic Data Systems Inc, Seaford DE) every two hours for 8 hours.

#### Oral glucose tolerance test (OGTT)

For OGTT we followed a previously described method [8] with minor modifications, as follows. We performed OGTT after 7 days of cold housing and Dox 600 mg/kg HFD diet. On day 8, we fasted mice for 5 h. We administered 2.5 g of glucose/kg of body weight by oral gavage and collected tail blood at the indicated time points for measurement of glucose. For glucose measurement, we used a Contour Next EZ glucometer (Bayer HealthCare LLC).

#### RNA extraction, qPCR and primers

For RNA extraction and qPCR, we followed the same method as previously described [8]. Briefly, for RNA extraction we homogenized ∼50 mg of tissue in 1 ml of QIAzol (Qiagen, Cat #79306) using a Tissue Lyser II and stainless-steel beads (Qiagen, 69989). After homogenization, we centrifuged the samples at 14,000 × *g* for 5 min and then removed the floating fat layer from the top by pipetting; we then added 200 ul chloroform and centrifuged the samples at 14,000 g for 15 min. We collected the supernatant and used an RNA extraction kit (Cat #74104, Qiagen) to obtain RNA. During RNA purification we used the RNase-Free DNase Set (Cat #79254, Qiagen) for DNA digestion. For qPCR we prepared cDNA with iScript kit, (Bio-Rad Cat # 1708891) using 1 μg of RNA and followed the manufacturer’s instructions, using the following cycles and temperatures: 5 min at 25 °C, 30 min at 42 °C, 5 min at 85 °C and hold at 4 °C. After the cDNA preparation, we performed qPCR using Power Sybr green (0.1 μM final concentration for primers) on Applied Biosystem’s Viaa7 machine. Comparative Ct method (ΔΔ Ct) was used to analyze all qPCR data. Expression was normalized to that of the 18S ribosomal subunit as endogenous control, and the relative expression was calculated in comparison with the reference sample that is indicated in each figure. Primers were designed using Primer Express 3.0.1 (Applied Biosystems) or Primer Blast National Center for Biotechnology Information. Primers used for qPCR were as follows: *Plin5*, forward GAGGCAGCAACAGGGCTACT, reverse CAAAGAGTGTTCATAGGCGAGATG; *18S forward* GAG CGA AAG CAT TTG CCA AG, reverse GGC ATC GTT TAT GGT CGG AA; *Ucp1* forward CCC TGG CAA AAA CAG AAG GA, reverse AGC TGA TTT GCC TCT GAA TGC; *Dio* forward AAG AAG CAC CGG AAC CAA GA, reverse GGC GGC AAG GAG AAA CG; *Elov3* forward GCC AAA CTG AAG CAT CCT AAT CTT, reverse CCC AGA ACC ATC TGC AGA ATC; *Ppargc* forward TGC CAT TGT TAA GAC CGA G, reverse TTG GGG TCA TTT GGT GAC.

#### Tissue protein lysate preparation, Western Blot and antibodies

For protein lysate preparation and Western blot we followed the same methods as previously described [8].We homogenized tissue (∼50 mg per sample) in RIPA buffer containing 50 mM tris(hydroxymethyl)aminomethane, 140 mM sodium chloride, 0.1% sodium dodecyl sulfate,1% triton X-100, 0.1% sodium deoxycholate, and 0.5 mM ethylene glycol-bis(2-aminoethylether)-*N,N,N*′*,N*′-tetraacetic acid, using a Tissue Lyser II with stainless-steel beads (Qiagen, Germany). After homogenization, we centrifuged the samples at 14,000 × *g* for 10 min at 4 °C to remove cell debris and collected the supernatants. We measured protein concentration using Pierce® 660 nm Protein assay reagent (Thermo Scientific, Cat # 22660). We mixed the samples with 2× protein sample loading buffer (62.5 mM Tris-HCL, 25% glycerol, 2% SDS, and 0.1% Orange G). For protein electrophoresis, we loaded 20 µg of protein per sample into premade gels [Criterion^TM^ TGX ^TM^ 4–20% (Bio-Rad, Cat # 5671094) or AnyKD (Bio-Rad, Cat # 5671124)] and for protein transfer, we used the Criterion Blotter System^TM^ with nitrocellulose membrane (Bio-Rad, Cat # 1620112). We blocked the membranes post-transfer with 5% nonfat dry milk diluted in Tris-buffered saline, pH 7.4, with 0.1% Tween-20 (TBS-T) for 1 h, and then incubated with the indicated primary antibody with 3% BSA diluted in TBS-T (BSA-TBS-T) for 12–16 h at 6 °C. After primary antibody incubation, we washed the membranes three times for 5 min each with TBS-T, and then incubated with the appropriate secondary antibody from Li-Cor in BSATBS-T for 30 min. After secondary antibody, we washed membranes three times for 5 min each in TBS-T. We visualized the immunoblotted proteins with the Odyssey CLx near-infrared imaging system (Li-Cor).

We used the following primary antibodies: PLIN5 (Progen Cat # GP31), GDI (Bickel Lab) [14], Histone H3 (Cell signaling Cat # 9717), Cox4 (Cell signaling Cat# 4850), TIM23 (Cell signaling Cat # 34822), TOM20 (Santa Cruz sc-11021), Actin (Cell Signaling Cat# 4967).

#### Glucose and fatty acid uptake

For glucose and fatty acid uptake we followed the methods as described previously [15] with minor modifications. We administered by oral gavage 1 mg/g body weight glucose with 20 µCi deoxy-D-glucose, 2-[1-^14^C]- (Perkin Elmer, Cat # NEC495A00) per mouse and ^3^H-triolein (Perkin Elmer, Cat # NET43100) as tracers. After 1-h, mice were sacrificed and tissues of interest were harvested and weighed, small pieces of each tissue were cut, weighed, and solubilized using Solvable^TM^ (0.1 ml per 10 mg of tissue) and 200 µl were added to 5 ml of scintillation cocktail in a glass scintillation vial. Radioactivity was measured using a scintillation counter (Beckman Coulter, LS6000).

#### Histology

We performed histology as previously described [8]. To obtain mouse tissue samples for histology, we performed cardiac perfusion under ketamine anesthesia. After cardiac perfusion with 4% paraformaldehyde in phosphate-buffered saline (PBS), pH 7.4, we dissected tissues and fixed with 4% paraformaldehyde solution overnight. The UTSW Molecular Pathology Core Facility performed paraffin embedding, sectioning, H&E staining, and Oil red O staining. We acquired bright-field images using a Leica DM 4000B microscope.

#### Electron microscopy

For electron microscopy we followed the methods as previously described [8]. For transmission electron microscopy sample collection, we performed cardiac perfusion under ketamine anesthesia with a perfusion buffer (4% paraformaldehyde, 1% glutaraldehyde, and 0.1 M sodium cacodylate, pH 7.4), and we dissected BAT into 1 mm pieces that were then fixed with 2.5% glutaraldehyde and 0.1 M sodium cacodylate, pH 7.4. Further processing of the samples was performed by the UTSW Electron Microscopy Core as follows: tissue samples were rinsed in 0.1 M sodium cacodylate buffer and postfixed in 1% osmium tetroxide and 0.8% potassium ferricyanide in 0.1 M sodium cacodylate buffer three times for 3 h at room temperature. After three rinses in water, they were stained en bloc with 4% uranyl acetate in 50% ethanol for 2 h. Next, the samples were dehydrated with increasing concentrations of ethanol, transitioned into resin with propylene oxide, infiltrated with Embed-812 resin, and polymerized in a 60 °C oven overnight. Blocks were sectioned with a diamond knife (Diatome) on a Leica Ultracut 7 ultramicrotome (Leica Microsystems), collected onto copper grids, and post stained with 2% aqueous uranyl acetate and lead citrate. Images were acquired on a JEOL 1400 Plus electron microscope and photographed with a BIOSPR16 camera.

#### Quantification of mitochondria and lipid droplet contact sites

We performed quantification of mitochondria and lipid droplet contact sites as previously described [8, 16]. For quantification of LD–mitochondria contact sites, we used Image J software NIH Ver 1.5. We quantified mitochondria in contact with LDs by count and contact area as % of mitochondrial perimeter or % of LD perimeter (*n* = 10 EM fields per genotype at 1000x magnification).

#### mtDNA quantification

We performed mtDNA quantification as previously described [8]. For isolation of cellular total DNA, 25 mg of tissue were homogenized in PBS (ph. 7.5) using a Tissue Lyser II and stainless-steel beads (Qiagen, 69989). Samples were processed with Qiagen’s QIAamp DNA Mini Kit (51304) per the kit instructions. Mitochondrial DNA was amplified using primers specific for the mitochondrial cytochrome *c* oxidase subunit 2 (COX2) gene and normalized to genomic DNA by amplification of the ribosomal protein s18 (rps18) nuclear gene, using quantitative PCR. We used primers that were previously described [17] as follows: *Rsp18* forward TGT GTT AGG GGA CTG GTG GAC A and reverse CAT CAC CCA CTT ACC CCC AAA A and *Cox2* forward ATA ACC GAG TCG TTC TGC CAA T and reverse TTT CAG AGC ATT GGC CAT AGA A.

#### Mitochondria isolation

We isolated crude and pure mitochondria as described previously [18]. Mice were euthanized (Control or BiKOPLIN5) by isoflurane overdose and neck dislocation. We dissected BAT and rinsed the tissue in cold buffer containing 225-mM mannitol, 75-mM sucrose, 0.5% BSA, 0.5-mM EGTA and 30-mM Tris–HCl pH 7.4.

We then minced 50-100 mg of tissue into approximately 1 mm pieces and resuspended the pieces in the same buffer used for rinsing the BAT (500 μl per each 100 mg of tissue). We then transferred BAT to a 10 ml glass/Teflon Potter Elvehjem homogenizer and homogenized the BAT pieces using a Teflon pestle with eight strokes at 1,500 r.p.m. We then we centrifuged the homogenate in a 15 ml polypropylene centrifugation tube at 740*g* for 5 min at 4 °C. We collected the supernatant and centrifuged one more time at 740*g* for 5 min at 4 °C. We collected the supernatant and centrifuged at 9,000*g* for 10 min at 4 °C. After this step we discarded the supernatant and carefully resuspended the pellet in 1 ml of buffer containing 225-mM mannitol, 75-mM sucrose, 0.5% BSA and 30-mM Tris–HCl pH 7.4. We then centrifuged the mitochondrial suspension at 10,000*g* for 10 min at 4 °C. This pellet was crude mitochondria. We resuspended the pellet in 50-200 μl of buffer containing 250-mM mannitol, 5-mM HEPES (pH 7.4) and 0.5-mM EGTA and then measured protein concentration. For subsequent mitochondrial respiration experiments we resuspended these crude mitochondria in resuspension buffer (225-mM mannitol, 75-mM sucrose, 0.5% BSA and 30-mM Tris–HCl pH 7.4) or for Western blotting we added 1X protein loading buffer (Li-Cor Cat # 928-40004).

For pure mitochondria isolation we used as starting material the crude mitochondria obtained as described above. First, we added 2 ml of buffer containing 225-mM mannitol, 25-mM HEPES (pH 7.4), 1-mM EGTA and 30% Percoll (vol/vol) to an ultracentrifuge tube, then carefully layered 200 μl of the crude mitochondria obtained above. Then we layered 1 ml of buffer containing 250-mM mannitol, 5-mM HEPES (pH 7.4) and 0.5-mM EGTA and centrifuged at 95,000*g* for 30 mins. After centrifugation we collected the pure mitochondria from the bottom of the tube using a Pasteur pipette. We resuspended the pellet in 2 ml of buffer containing 250-mM mannitol, 5-mM HEPES (pH 7.4) and 0.5-mM EGTA and centrifuged at 6,300*g* for 10 minutes. We repeated this step one additional time and discarded the supernatant. Finally, we resuspended the pellet in 100 μl of buffer containing 250-mM mannitol, 5-mM HEPES (pH 7.4) and 0.5-mM EGTA. We measured protein concentration, and the mitochondria were used for Western blotting or for protease protection assays.

#### Protease protection assay

We performed protease protection assays using pure mitochondria obtained as described above. To the resuspended pellet of pure mitochondria (in buffer containing 250 mM mannitol, 5 mM HEPES, pH 7.4, and 0.5 mM EGTA), we added Proteinase K (100 µg/ml) and incubated the samples on ice for 15 min. We stopped protease digestion by adding Phenylmethylsulfonyl Fluoride PMSF (2 mM) and incubating the samples on ice for 5 min. We measured protein concentration and further processed the sample for Western blot with 1X protein loading buffer (Li-Cor Cat # 928-40004).

#### Mitochondria respiration

For assessment of mitochondrial respiration, we used crude mitochondria isolated from intrascapular BAT of BiKOPLIN5 mice as described above. To measure oxygen consumption rate, we used the NeoFox apparatus as follows. We suspended mitochondria corresponding to 50 μg of mitochondrial protein in a buffer containing 250-mM mannitol, 5-mM HEPES (pH 7.4) and 0.5-mM EGTA. Oxygen consumption rate (OCR) was determined before and after sequential injections of the following compounds pyruvate (5mM), Guanosine 5′-diphosphate sodium (GDP-1 mM), Adenosine 5′-diphosphate sodium (ADP-450 μM), oligomycin (2 μg/ml) and Carbonyl cyanide 4-(trifluoromethoxy) phenylhydrazone (FCCP-1 μM). We calculated OCR as μmol/l/min/μg of protein.

#### Software

For WB band intensity analysis, we used Image Studio Ver. 3.1 (Licor Biosciences). For qPCR Ct values analysis, we used Quant Studio Real-time qPCR software Ver. 3.1 (Applied Biosystems). For colorimetric microplate assays (protein quantification) we used Gen5 Ver 2.01.14 software (Bio Tek Instruments, Inc.). For quantification of LD–mitochondria contact sites, we used Image J software NIH Ver 1.53.

#### Statistical analysis

For statistical analysis we used GraphPad Prism version 10.0.1 for MacOS, GraphPad Software, La Joya California USA, www.graphpad.com.

For all the experiments, data are representative of at least three independent experiments and all attempts to reproduce were successful. P values are indicated in figures or figure legends. Statistical analyses were performed using Student’s t test if two groups were analyzed or ANOVA followed by Tukey posttest if more than two groups were analyzed. Statistical significance is defined as p<0.05.

## Results

### Creation and validation of an inducible mouse model of BAT-specific PLIN5 knockout

To create this doxycycline-inducible knockout mouse line, we crossed our previously described *Plin5^loxp/loxp^* mouse strain, in which we had introduced LoxP sites flanking exons 3 through 8 of the *Plin5* gene by homologous recombination in C57BL/6 embryonic stem (ES) cells [8], with transgenic mice expressing the “tet-on” transcription factor rtTA under the control of the *Ucp1* gene promoter (*Ucp1-*rtTA), which was generously provided by Phillip Scherer [12]. Finally, we crossed *Plin5 ^loxp/loxp^; Ucp1*-rtTA mice with mice carrying a tetracycline-responsive Cre recombinase (*TRE*-Cre) [13] to produce BiKOPLIN5 mice. Littermate control (Control) mice lacked the TRE-Cre allele (Fig. 1a).

**Figure 1.**
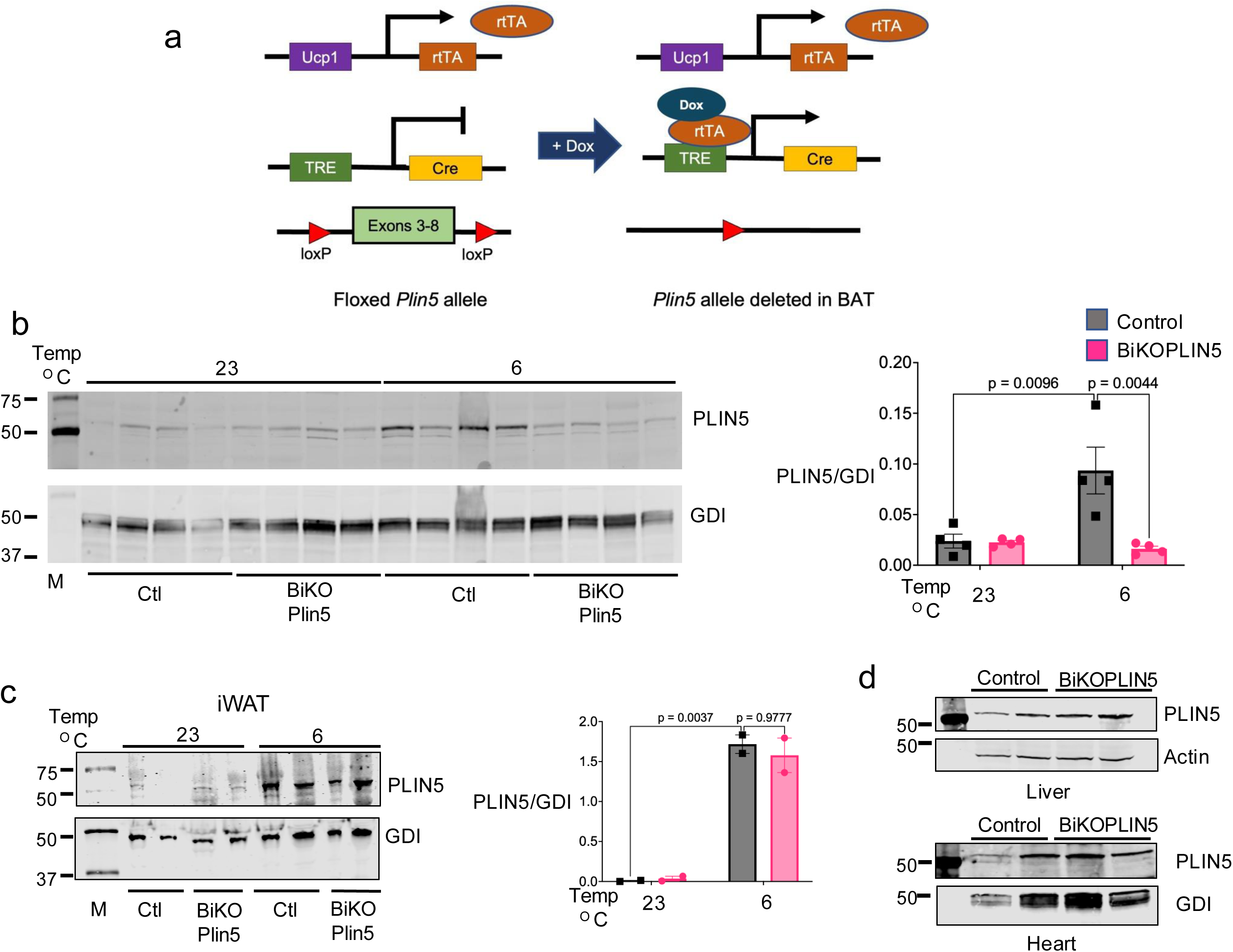
Validation and generation of BiKOPLIN5 mice. (a) Schematic representation for the strategy for doxycycline-inducible disruption of *Plin5* gene. (b) Western blot (WB) depicting PLIN5 in BAT (left panel) and quantification (right panel) and (c) iWAT (left panel) and quantification (right panel) in BiKOPLIN5 and control mice housed at 23 °C or 6 °C for 7 days. (d) WB depicting PLIN5 from liver (left panel) and heart (right panel) of Control or BiKOPLIN5 mice. For WB experiments, mice were housed at the indicated temperatures and administered Dox for 7 days.”

We evaluated different doxycycline doses, durations of doxycycline treatment, and housing temperatures to determine optimal conditions to achieve significant knockout of PLIN5 expression specifically in BAT without activating compensatory beiging in iWAT. As expected, based on our previously published data that PLIN5 is induced in BAT of C57BL/6J mice during housing at 6 °C [8], PLIN5 was increased 4-fold in the BAT of Control mice after 7 days of doxycycline diet and housing at 6 °C (Fig. 1b). In contrast, under the same conditions, PLIN5 in the BAT of BiKOPLIN5 mice was ∼80% less than in that of Control mice. No significant differences in PLIN5 protein in iWAT, liver, or heart were observed between BiKOPLIN5 and Control mice (Fig. 1c, d). Seven days of doxycycline treatment of mice housed at 23 °C was not associated with reduced PLIN5 in BAT or iWAT of either BiKOPLIN5 or Control mice.

### Inducible *Plin5* BAT knockout causes reduction on thermogenic gene expression in BAT

To study the effects of acute BAT *Plin5* knockout on thermogenic gene expression in BAT and iWAT, we performed real-time quantitative polymerase chain reaction (qPCR) on RNA isolated from BAT and iWAT after 7 or 21 days of Dox administration and housing at 23 ^°^C or 6 ^°^C. Whereas *Plin5* mRNA increased in Control mice after 7 days at 6 ^°^C compared with those housed at 23 ^°^C, *Plin5* mRNA in BiKOPLIN5 mice did not change significantly (Figure 2a). Similarly, in the BAT of BiKOPLIN5 mice, expression of most thermogenic genes (*Ucp1*, *Dio*, and *Ppargc*) after 7 days at 6 ^°^C was significantly lower than in Control mice, but there were no differences between these genotypes in iWAT (Fig. 2b). After 21 days of Dox diet, we observed much reduced *Plin5* expression in the BAT of BiKOPLIN5 at both 23 ^°^C and 6 ^°^C compared with Control mice, as well as in iWAT at 6 ^°^C. At the three-week time point, expression levels of *Ucp1*, *Dio*, *Elov3* and *Ppargc* in BAT were reduced in iWAT, but only that of *Ppargc* was statistically significant (Fig. 2c, d).

**Figure 2.**
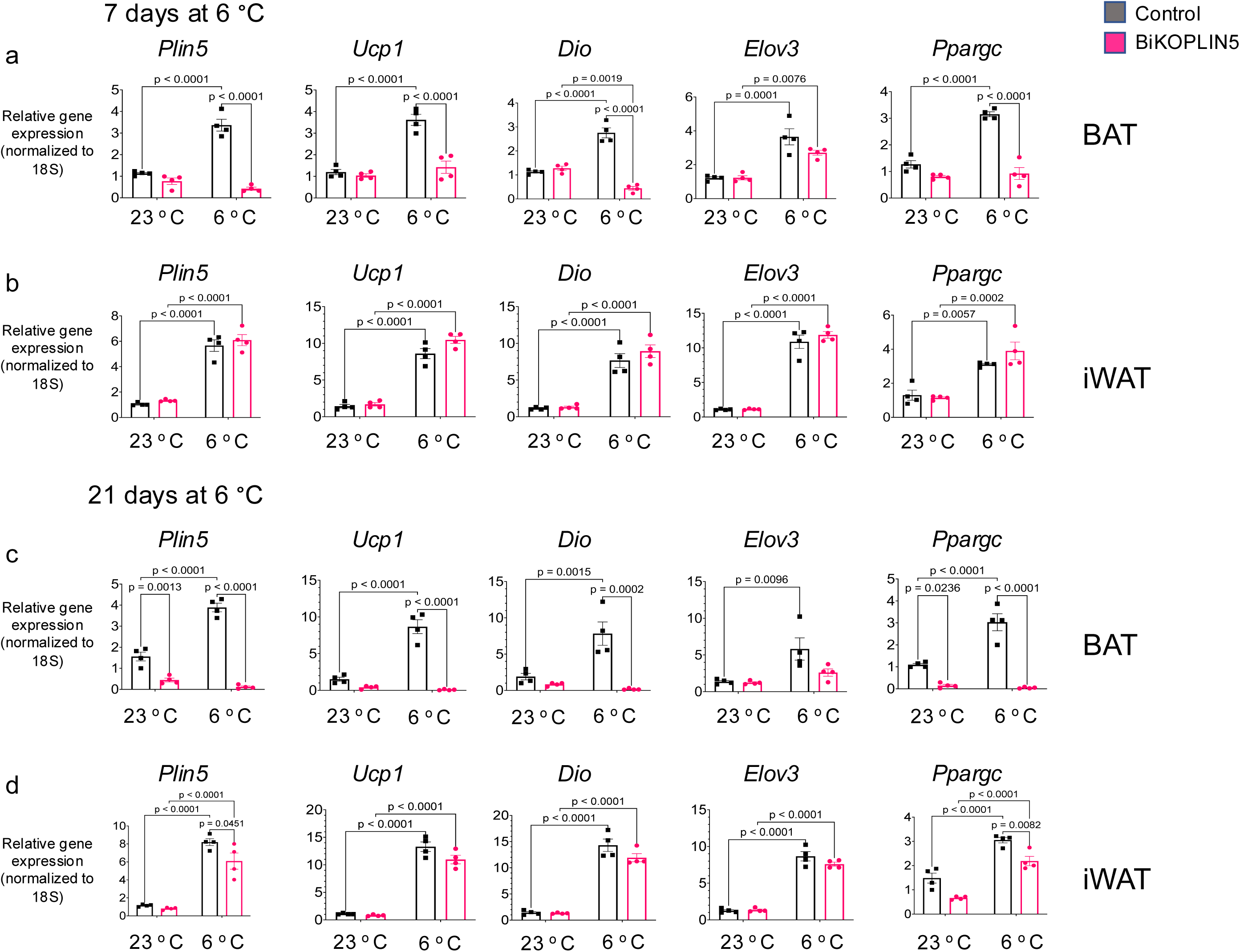
BiKOPlin5 mice exhibit reduction in thermogenic gene expression in BAT. (a) Quantitative polymerase chain reaction (qPCR) for the indicated genes from BAT or (b) iWAT from Control or BiKOPlin5 mice housed at 6 °C for 7 days. (c) qPCR for the indicated genes from BAT or (d) iWAT from control or BiKOPlin5 mice housed at 6 °C for 21days. (n=4 per group). Statistical analysis was performed using two-way ANOVA followed by Tukey post-test for multiple comparisons. Significant p values (p < 0.05) are shown in the figure.

To study the effects of acute *Plin5* gene disruption specifically in BAT but sparing iWAT, subsequent experiments were conducted on day 8 following 7 days of housing the experimental mice at 6 ^°^C and of Dox-chow diet.

### Acute BAT PLIN5 knockout causes cold intolerance but not glucose intolerance

First, we evaluated body weight and daily food intake when mice were housed at 23 °C without Dox diet and then during 7 days of cold exposure with Dox diet. Food intake was calculated as the average of the 3 days before day 0, when mice were maintained at 23 °C and not yet on the Dox diet, and again as the average of days 5, 6, and 7 of cold exposure and Dox diet. Body weight was measured at day 0 and again at day 7. We found no differences in food intake or body weight between Control and BATiKOPLIN5 mice either before or after cold exposure and Dox diet (Figure 3 a, b). To determine the effect of acute PLIN5 deficiency in BAT on glucose tolerance in the context of high-fat diet (HFD) induced obesity, we fed male or female mice with 60% HFD without Dox for 8 weeks and housed the mice at 23 ^°^C. After 8 weeks we shifted the mice to housing at 6 ^°^C and initiated Dox diet with 60% HFD (HFD-Dox) for 7 days. On day 8-day we performed OGTT as described in Methods. Glucose tolerance was similar between control and BiKOPLIN5 mice in both male (Figure 3c) and female mice (Figure 3d).

**Figure 3.**
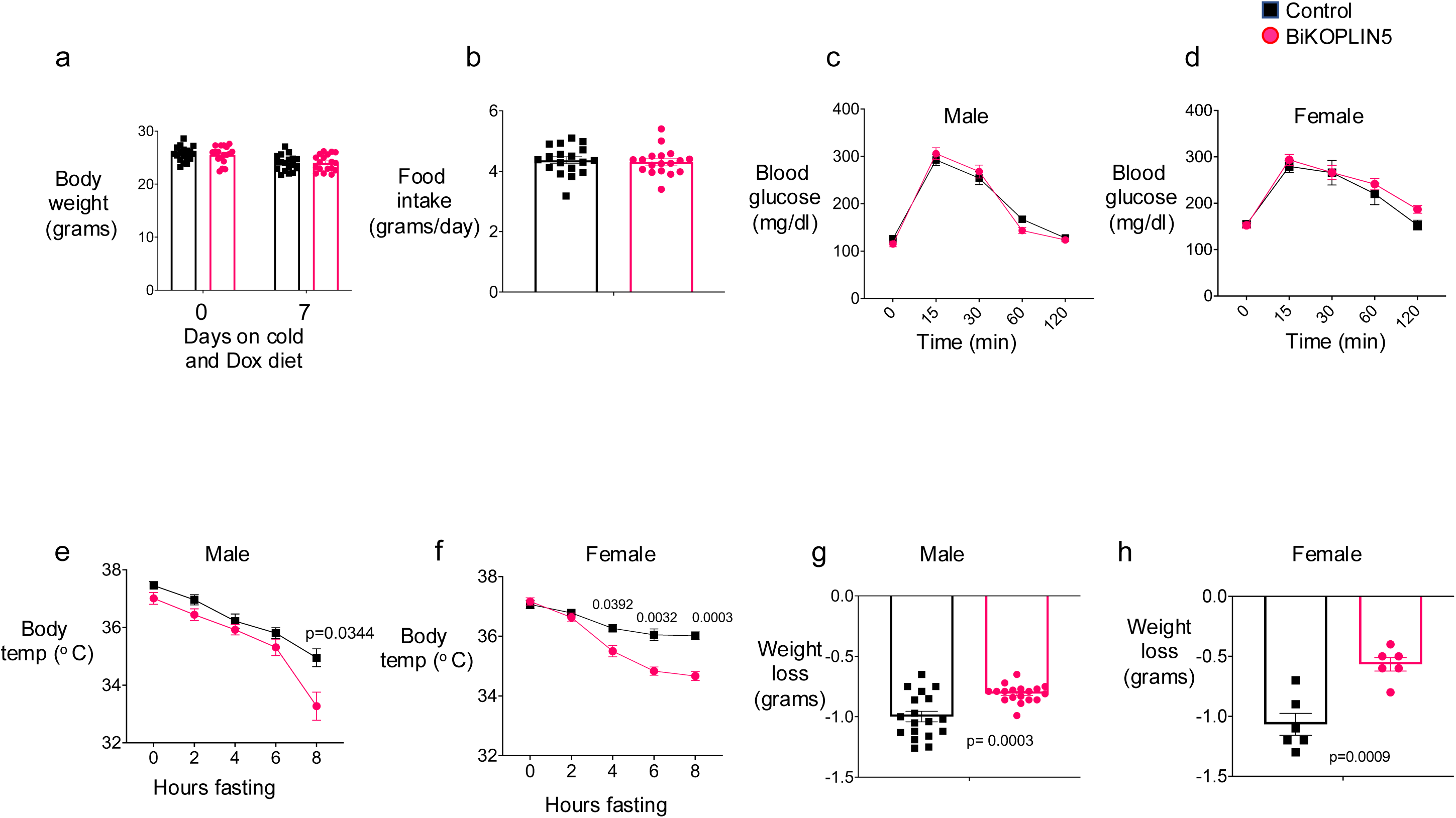
BiKOPLIN5 showed reduced cold tolerance but not glucose intolerance. (a) Body weight and (b) food intake of Control and BiKOPLIN5 mice fed a chow diet containing 600 mg/kg Dox and exposed to cold for 7 days (n = 18 per group). (c) Oral glucose tolerance tests (OGTT) in males and (d) females Control and BiKOPLIN5 mice. For OGTT studies, mice were fed a 60% HFD and housed at 23 °C for 8 weeks, then switched to a 60% HFD containing 600 mg/kg Dox and housed at 6 °C. OGTT was performed on day 8 of Dox treatment and cold exposure (n = 10–14 per group). (e) Cold tolerance tests in male and (f) female Control and BiKOPLIN5 mice. (g) Body weight changes during the cold tolerance test in males and (h) females. For cold tolerance assays, n = 18–19 for males per group and n = 6 for females per group. Statistical analysis was performed using an unpaired Student’s *t*-test. Significant *p*-values (*p* < 0.05) are indicated in the figure.

Given the reduction in thermogenic gene expression (Figure 2a) that we observed in BiKOPLIN5 mice, we performed a cold tolerance test to assess whether acute PLIN5 deficiency in BAT results in cold intolerance. First, we implanted a temperature transponder subcutaneously and allowed the mice to recover for 3 days. Then we initiated Chow-Dox diet and housed the mice at 6 ^°^C for 7 days. On day 8 we fasted the mice starting at 8 a.m. and measured body temperature every two hours for 8 hours. Male BiKOPLIN5 mice showed consistently lower body temperatures than Control mice at every time point with statistically significant changes at 8 hours cold exposure (Figure 3e). Female BiKOPLIN5 mice had statistically significant lower body temperatures than female Control mice at hours 4, 6, and 8 of the test. (Figure 3f). We also measured body weight before and after fasting during the cold tolerance test. Both male and female BiKOPLIN5 mice lost less weight during the test than Control mice (Figure 3g-h).

### Acute BAT *Plin5* knockout decreases BAT tissue fatty acid uptake

BAT augments triglyceride and fatty acid clearance especially during cold exposure [15] and previously, we found that BAT PLIN5 overexpression increases BAT fatty acid uptake but has no effect on glucose uptake by that tissue [8]. To evaluate if acute induction of PLIN5 deficiency in BAT changes fatty acid or glucose uptake, we performed a combined oral triacylglycerol and glucose tolerance test using H^3^labeled triolein and C^14^ labeled glucose, as previously described [15], in BiKOPLIN5 and Control mice housed for 7 days at 6 ^°^C and fed Chow-Dox diet One hour after administration of the labeled substrates via oral gavage, we harvested the indicated tissues and assessed uptake of the labeled substrates by scintigraphy, as described in Methods. Fatty acid uptake was decreased in the BAT of BiKOPLIN5 mice compared with Control mice, but there were no statistically significant differences in iWAT, gonadal WAT (gWAT), heart, or liver (Figure 4a). There also were no differences in glucose uptake between BiKOPLIN5 and Control mice in any of these tissues, including BAT (Figure 4b).

**Figure 4.**
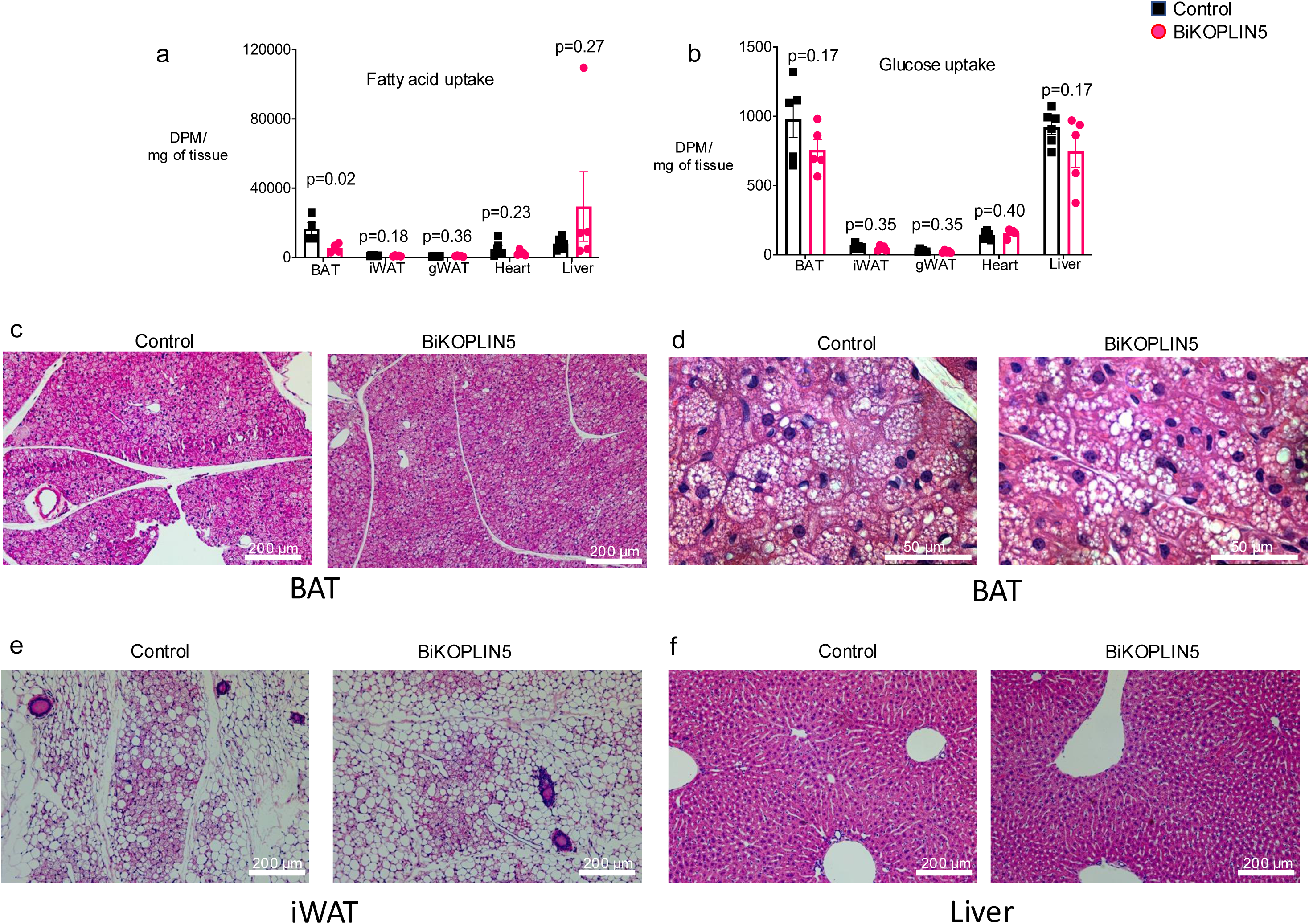
BiKOPLIN5 showed reduced fatty acid uptake but not glucose uptake in BAT. (a) Fatty acid and (b) glucose uptake on the indicated tissues from BiKOPLIN5 or Control mice. For this assay the mice were fed 600 mg/kg diet chow Dox and housed at 6 °C for 7 days. Assay was performed on day 8. n= 6 per group. Statistical analysis was performed using unpaired Student *t* test. Significant p values (p < 0.05) are shown in the figure. (c) Representative image of hematoxylin and eosin staining from BAT at 10x magnification or (d) 40x magnification from control or BiKOPLIN5 mice. (e) Representative image of hematoxylin and eosin staining from iWAT at 10x magnification from control or BiKOPLIN5 mice. (f) Representative image of hematoxylin and eosin staining from liver at 10x magnification from control or BiKOPLIN5 mice. Scale bar is shown in the figure. n=3 per group.

In an independent cohort of BiKOPLIN5 and the Control mice, we studied the histology of the BAT, iWAT, and livers. By H&E staining we found no differences in BAT (Figure 4c and d), iWAT (Figure 4e) or liver (Figure 4f) between the Control and BiKOPLIN5 mice. Both genotypes showed extensive browning of iWAT likely due to the 7 days of cold exposure we used to promote expression of Cre-recombinase in our mouse model (Figure 4e). The observed browning was consistent with the induction of thermogenic gene expression we measured in iWAT (Figure 2b). However, these changes in gene expression and histology in iWAT were not sufficient to overcome PLIN5 deficiency in BAT for the purpose of cold tolerance.

### BAT Plin5 knockout impairs mitochondrial morphology and respiration

Given the cold intolerance we observed in mice with acute PLIN5 deficiency, we next evaluated mitochondria morphology and function in the BiKOPLIN5 mice by transmission electron microscopy of BAT tissue and respirometry of isolated mitochondria. Similar to constitutive PLIN5 KO mice [8], brown adipocytes in cold-stressed BiKOPLIN5 mice had dysmorphic mitochondria with loss of cristae and swelling of mitochondria across the whole tissue (Figure 5a-top panels, lower magnification). These differences were more clearly observed at higher magnification (Figure 5a, bottom panels). Similar to the constitutive PLIN5 knockout mice from our previous work [8], BiKOPLIN5 did not show changes in mitochondria/lipid droplet contact sites (Figure 5b) in terms of the percentage of mitochondria in contact with lipid droplets, the ratio of mitochondria-lipid droplet contact surface to mitochondrial perimeter, and the ratio of mitochondria-lipid droplet contact surface to lipid droplet perimeter. BiKOPLIN5 mice also showed a trend to reduced mtDNA content, but this was not statistically significant (Figure 5c). Additionally, we tested mitochondrial function by measuring oxygen consumption rate (OCR) for mitochondrial isolated from BAT of Control or BiKOPLIN5 mice housed at 6 ^°^C and fed with Dox for 7 days. The BiKOPLIN5 mitochondria in comparison to Control mitochondria exhibited a blunted increase in mitochondrial OCR after the administration of pyruvate, as well a significant reduction in maximal uncoupled respiration following the addition of carbonyl cyanide 4(trifluoromethoxy)phenylhydrazone (FCCP) (Figure 5d). After addition of guanosine diphosphate (GDP), an inhibitor of UCP1mediated uncoupled respiration, the rate of decline in oxygen consumption was nearly flat in BiKOPLIN5 mitochondrial compared with a steep decline in Control mitochondria.

**Figure 5.**
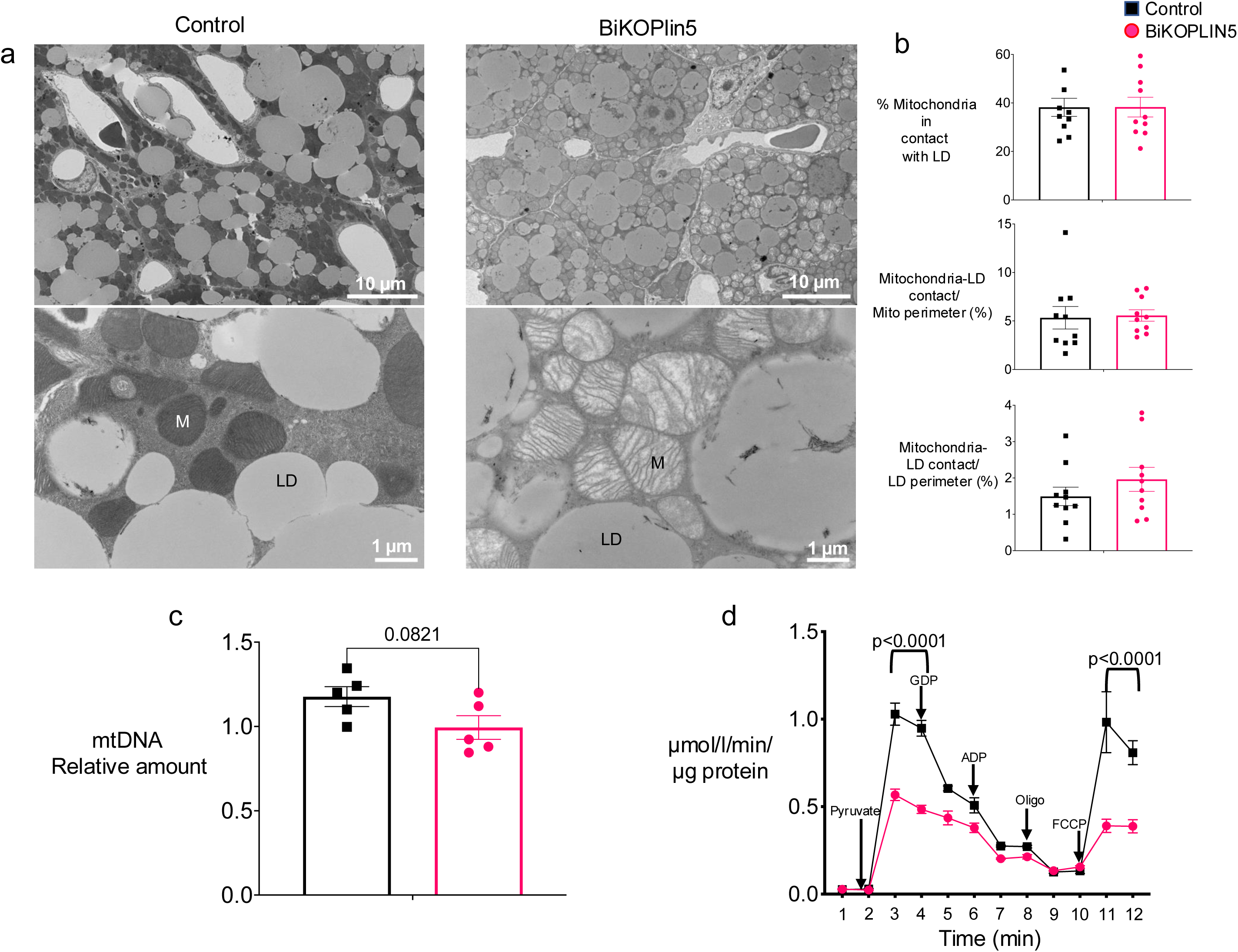
Plin5 deletion in BAT causes mitochondrial dysfunction. (a) BAT electron micrograph from Control (left panel) or BIKOPLIN5 (right panel) mice housed at 6 °C and fed with 600 mg/kg Dox diet for 7 days. Top panel is 800x and lower panel is 5000x magnification. M=mitochondria, LD=lipid droplets. (b) Electron micrograph quantification of mitochondria in contact with lipid droplets (top panel), mitochondria-lipid droplet contact/mitochondria perimeter (middle panel) and mitochondria lipid droplet contact/lipid droplet perimeter (bottom panel). n=10 EM fields per group. Statistical analysis was performed using unpaired Student *t* test. Significant p values (p < 0.05) are shown in the figure. (c) BAT Mitochondrial DNA quantification from control or BiKOPLIN5 mice housed at 6 °C and fed with 600 mg/kg Dox diet for 7 days (d) Oxygen consumption rate (OCR) from BAT mitochondria isolated from control or BiKOPLIN5 mice housed at 6 °C and fed with 600 mg/kg Dox diet for 7 days. Mitochondria were sequentially injected with Pyruvate, GDP, ADP, Oligomycin and FCCP n=3 per group (d). Statistical analysis was performed using unpaired Student *t* test. Significant p values (p < 0.05) are shown in the figure.

### PLIN5 localizes to the outer mitochondrial membrane in brown adipocytes during cold exposure of mice

It has been reported that PLIN5 localizes to mitochondria in skeletal muscle, as well in several cell lines [6, 7, 19]. However, whether PLIN5 localizes to mitochondria in brown adipocytes is not known. Based on the profound changes in mitochondrial morphology and function in the BAT of BiKOPLIN5 mice, we next tested whether PLIN5 localizes to the mitochondria of brown adipocytes.

To this end, we isolated pure mitochondria as described in Methods from the BAT of C57BL6/J male mice that had been housed at either 23 °C or 6 °C for 48 h. The isolated mitochondria were assessed by a protease protection assay with Proteinase K to determine whether PLIN5 localizes to the outer mitochondrial membrane (OMM) or the inner mitochondrial membrane (IMM). We used antibodies to proteins that reside specifically in each of these compartments to assess localization of PLIN5 and of marker proteins: mitochondrial import receptor subunit TOM20 homolog (TOM20) as a marker of the OMM, mitochondrial import inner membrane translocase subunit Tim23 (TIM23) as a marker of the IMM. PLIN5 was only detected in the pure mitochondria fraction isolated from the BAT of mice following cold exposure and not from that of mice at room temperature (Figure 6a). Nor was PLIN5 detectable in the mitochondria isolated from BAT of BKOPLIN5 knockout mice (negative control). After adding proteinase K to the pure mitochondria sample, PLIN5 was no longer detectable by immunoblotting, as was the case for TOM20. On the other hand, the inner mitochondrial membrane marker TIM23 was resistant to proteolysis by proteinase K. These data are consistent with PLIN5 localization to the OMM (Figure 6a).

**Figure 6.**
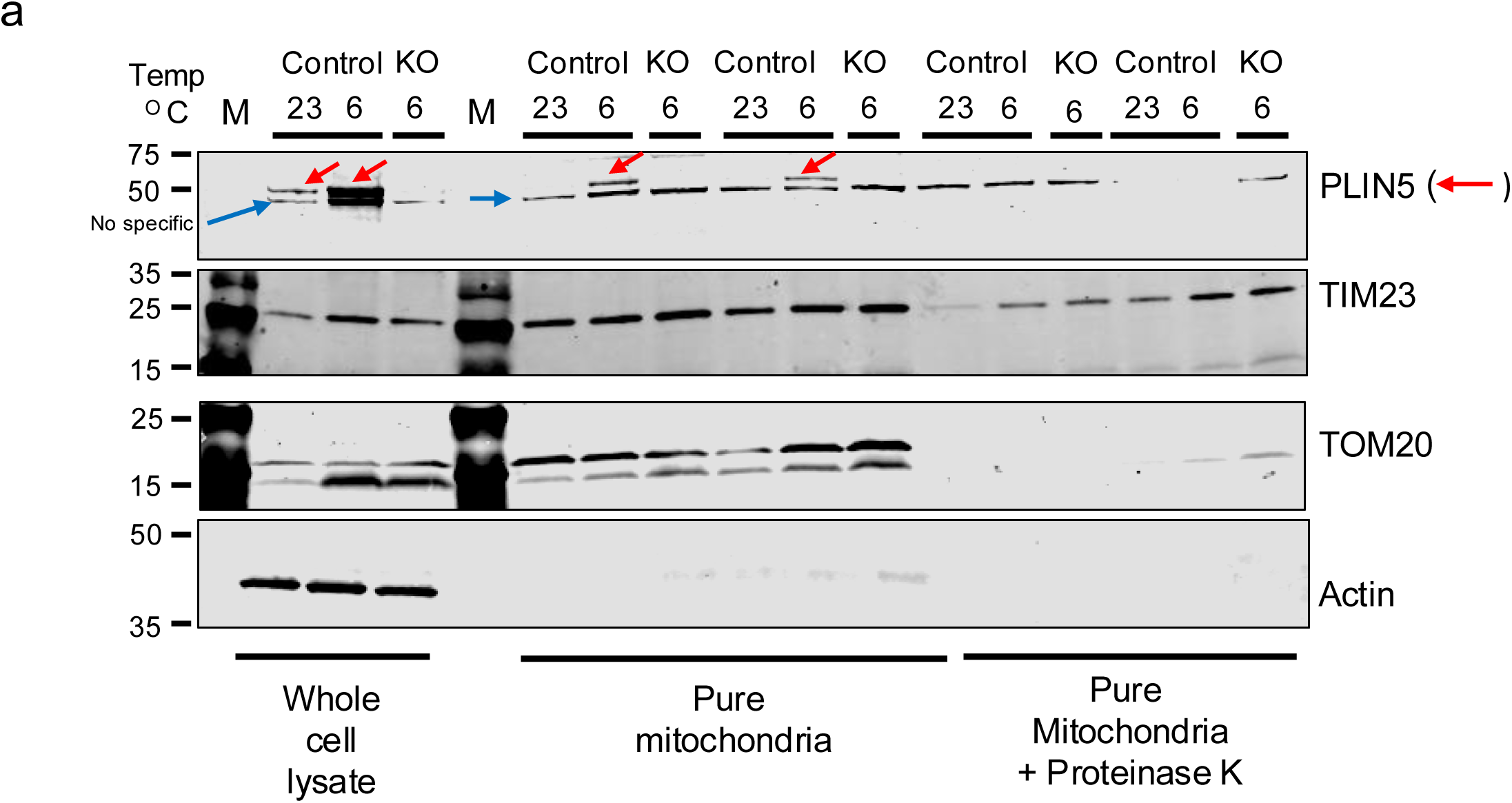
PLIN5 localize on the outer mitochondrial membrane in BAT. (a) Western blot for PLIN5, Actin, TIM23 and TOM20 from whole cell lysate and pure mitochondria with and without treatment with Proteinase K from control or BKOPLIN5 mice housed at 23 or 6 °C for 16 hours.

**Figure 7.**
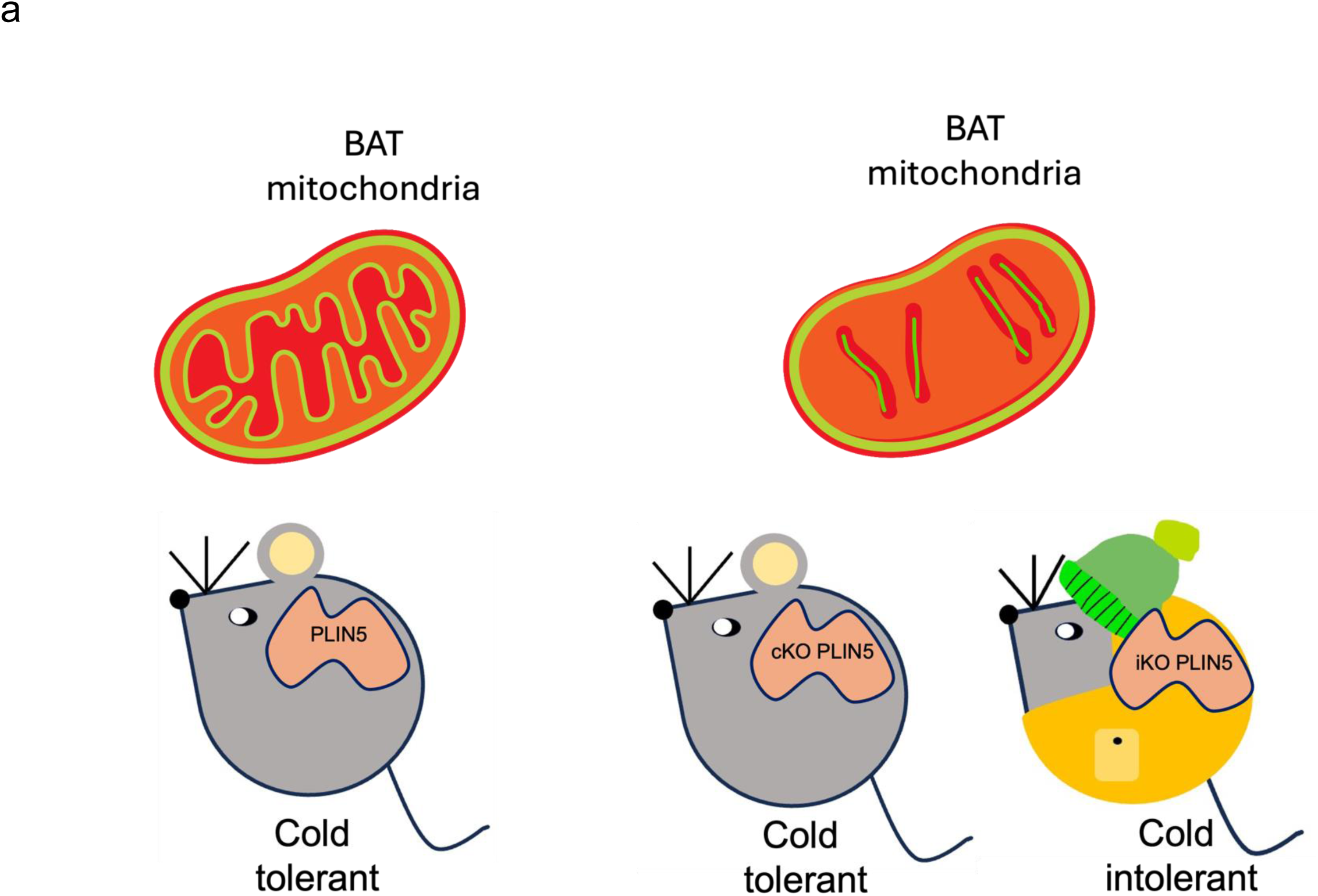
Working model of Perilipin 5 acute and chronic knockout in BAT. (a) Both acute and chronic PLIN5 loss led to impaired mitochondrial structure and function in BAT. However, the physiological consequences differ depending on the duration of PLIN5 depletion. In the acute setting, mitochondrial damage occurs before compensatory mechanisms can be engaged, resulting in pronounced cold intolerance. In contrast, chronic PLIN5 knockout also causes mitochondrial dysfunction but allows time for compensatory thermogenic activation in iWAT, which maintains thermogenesis and prevents overt cold intolerance. PLIN5-Control mice, cKOPLIN5-Constitutively KO mice (BKOPLIN5), iKOPLIN5-inducible KO mice (BiKOPLIN5)

## Discussion

Herein, we studied the effects of acute (7 days) depletion of PLIN5 in BAT of C57BL6/J mice using a doxycycline-inducible knockout of a floxed *Plin5* allele (UCP1*^rtTa^*; TRE-Cre; *Plin5^flox/flox^*). Previously, we demonstrated that constitutive, BAT-specific *Plin5* knockout (UCP1-Cre; *Plin5^flox/flox^*) in mice results in mitochondrial damage during cold exposure and reduced mitochondrial oxygen consumption rate [8], yet this chronic model did not exhibit differences in cold tolerance. With the additional evidence from the BiKOPLIN5 model presented in this report, we find that acute PLIN5 loss in brown adipocytes is associated with cold-induced mitochondrial damage and dysfunction, similar to the model of chronic PLIN5 deficiency in BAT, but with cold intolerance. Acute deficiency of PLIN5 in BAT resulted in significant BAT phenotypes, including reduced expression of thermogenic genes, reduced fatty acid uptake, reduced mitochondrial DNA content, decreased mitochondrial cristae density, swollen mitochondria, and impaired mitochondrial respiration, which together were associated with cold intolerance at 7 days. In this acute setting, there was insufficient time for browning of iWAT to the degree that would compensate for BAT dysfunction, at least at the level of thermogenic gene expression. A similar phenomenon was reported by Pereira et al., who reported that BAT-specific OPA1 knockout caused mitochondrial dysfunction but also compensatory WAT browning, which increased cold tolerance above even that of control mice [20]. Our two different models of BAT PLIN5 deficiency highlight the value of assessing genetic loss-of-function across both acute and chronic time frames, which is achievable through inducible gene knockout technology.

Though PLIN5 localization to mitochondria has been reported previously, our data in this study are the first to demonstrate in mouse BAT that endogenous PLIN5 enriches on the mitochondrial outer membrane in pure mitochondrial fractions during cold exposure. Carole Sztalryd’s lab first reported PLIN5 mitochondrial localization in 2011 by demonstrating that endogenous PLIN5, but not PLIN2 or PLIN3, co-fractionates with crude mitochondria isolated from rat left heart ventricle and that fluorescently tagged PLIN5 localizes to mitochondria in AML12 hepatocytes and HL1 cardiomyocytes [6]. The authors proposed that PLIN5 promotes lipid droplet–mitochondria contacts to facilitate fatty acid transfer for oxidation in mitochondria. More recently, Sarah Cohen’s group reported that interactions between the C-terminal 38 amino acids of PLIN5 and mitochondrial FATP4 promote lipid droplet-mitochondrial tethering for augmentation of fatty acid transport from lipid droplets to mitochondria for fatty acid oxidation [21].

Guenter Haemmerle’s group further reported that, in AML12 hepatocytes expressing fluorescently tagged PLIN5, mitochondria are tightly associated with PLIN5-coated lipid droplets, and that this interaction persists following PKA activation [19]. Moreover, his group demonstrated that expression of PLIN5 lacking its last three amino acids did not promote lipid droplet-mitochondria contacts in HEK-293T cells [19].

In contrast to these previous reports, we found no differences in lipid droplet–mitochondria contacts in BAT from constitutive PLIN5-KO mice, which suggests that PLIN5 is not required for these associations in BAT [8]. Our current study of acute PLIN5 deficiency confirms that PLIN5 is dispensable for lipid droplet contacts with mitochondria in BAT.

Two other groups have examined the role of PLIN5 in mitochondria–lipid droplet contact sites in thermogenic cells. The Orian Shirihai group reported that, compared with thermoneutrality, cold exposure of mice reduces LD–mitochondria interactions in brown adipocytes. They found that cytoplasmic mitochondria not associated with lipid droplets exhibit a greater capacity for fatty acid oxidation than peri-droplet mitochondria. Consistent with this interpretation, they showed that overexpression of full-length PLIN5 in brown adipocytes promotes LD–mitochondria association, whereas expression of a truncated PLIN5 lacking the last 20 amino acids fails to recruit mitochondria to lipid droplets [16]. In line with these observations, we previously found that cold exposure (6 °C vs. 23 °C) decreased LD–mitochondria contacts in BAT in but not in BATiPLIN5 mice (PLIN5 overexpression in BAT). Notably, in mice housed at 23 °C, BATiPLIN5 mice exhibited a reduction in LD–mitochondria contact number and contact area compared with Control mice [8]. Taken together, and in the context of the Shirihai group’s finding that cytoplasmic mitochondria in BAT are more oxidative than peri-droplet mitochondria, our previous published data suggest that PLIN5 overexpression in BAT results in a higher proportion of mitochondria with an oxidative phenotype relative to Control mice.

In contrast to these PLIN5 overexpression models, we did not observe differences in mitochondria–lipid droplet contact sites in BAT by electron microscopy quantification in either our constitutive BAT-specific PLIN5 knockout mice (Supplementary Figure 19 [8]) or the inducible BATiKOPLIN5 mice described in the current study. These findings suggest that, in BAT, PLIN5 is not required for the formation or maintenance of lipid droplet–mitochondria contact sites. The group of Carles Cantó has identified Perilipin 1 (PLIN1)—through its interaction with Mitofusin 2 (Mfn2)— as a mediator of lipid droplet–mitochondria tethering in BAT [22], Whereas PLIN5 may promote mitochondrial tethering to lipid droplets in many cell types, it is possible that this function is assumed by PLIN1 or a different protein in brown adipocytes. However, the roles of PLIN5 and PLIN1 in mitochondria contacts with lipid droplets is far from settled as the Pingsheng Liu group reported that mitochondria are tightly associated with lipid droplets in oxidative tissues but that their association is not dependent on either PLIN5 or PLIN1. This conclusion was based on the proteomic identification of equivalent levels of mitochondrial proteins in lipid droplet fractions isolated from the BAT of wildtype, PLIN5 deficient, and PLIN1 knockout mice [23]. Our findings support the notion that while PLIN5 overexpression may modulate the distribution or metabolic phenotype of mitochondrial subpopulations, PLIN5 itself is likely dispensable for basal droplet–mitochondria tethering in BAT. It may be that the focus of investigators on PLIN5 playing a role in lipid droplet-mitochondria contacts reflects the original discovery of PLIN5 as a member of the Perilipin family of lipid droplet proteins [1–3]. It is important to recall that PLIN5 also exists in a cytoplasmic pool and can move on and off the lipid droplet. We cannot rule out the possibility that cytoplasmic PLIN5, not lipid droplet-bound PLIN5, is the active player on the mitochondrial outer membrane [5, 24, 25].

If PLIN5 does not localize to mitochondria to promote the physical coupling of lipid droplets with mitochondria in BAT, then what is the purpose of this localization that is promoted by cold exposure of mice? We observed profound disruption of mitochondrial cristae structure and impaired mitochondrial respiration in the BAT of BiKOPLIN5 mice housed at 6°C, which was fully consistent with our findings in our previous report of constitutive PLIN5 knockout in BAT. While the precise mitochondrial function of PLIN5 in BAT mitochondria remains to be defined, we propose that during cold exposure PLIN5 may interact with outer mitochondrial membrane proteins to help preserve mitochondrial membrane integrity during the metabolic stress of increased fatty acid flux and high beta-adrenergic signaling. In this regard, Guenter Haemmerle’s group reported that full-length PLIN5 interacts with mitochondrial protein complexes involved in oxidative phosphorylation and mitochondrial dynamics in AC16 cardiomyocytes, whereas cardiomyocytes expressing a truncated PLIN5 variant lacking the final three amino acids fails to exhibit these interactions and does not promote lipid droplet-mitochondrial contacts. [19]. While these findings shed light on potential mechanisms for PLIN5 role in cardiomyocyte mitochondria, they may not elucidate its role in BAT, in which mitochondria are unique and function differently. Mitochondria in cardiomyocytes are optimized for efficient ATP production [26]; in contrast, mitochondria in BAT are specialized for thermogenesis at the expense of ATP production [27]. In BAT, UCP1-mediated uncoupled respiration and additional UCP1-independent thermogenic pathways play a central role in thermogenesis, which highlights the need to define PLIN5’s specific mitochondrial function in this tissue context [27]. Future work will focus on defining this mitochondrial role of PLIN5 in brown adipocytes.

In summary, together with our previous discovery that, upon its phosphorylation by PKA, PLIN5 translocate to the nucleus to activate the SIRT1–PGC-1α pathway [5], our data establish PLIN5 as a key integrator of lipid droplet function with thermogenic gene programs and mitochondrial function. While studies in heart, liver, and muscle have shown that PLIN5 coordinates fatty acid handling with oxidative metabolism, here we extend this concept to BAT by demonstrating that PLIN5 is enriched in mitochondria during adrenergic stimulation and is required to preserve mitochondrial structure and respiration during cold exposure. This mitochondrial role in BAT is distinct from the previously proposed function of PLIN5 in promoting lipid droplet–mitochondria contacts in other tissues. Instead, our findings support a model in which PLIN5 couples lipid mobilization with nuclear transcriptional responses and mitochondrial performance to meet the energetic demands of thermogenesis.

## Data availability statement

All data supporting the conclusions of this study are provided in the manuscript or may be requested from the corresponding author.

## Acknowledgments

We thank Philipp Scherer and UT Southwestern Touchstone Diabetes Center for the UCP1rtTA mouse. We thank the UT Southwestern Metabolic Phenotyping Core (Ruth Gordillo and Syann Lee), Electron Microscopy Core (Kate Luby-Phelps), Histo Pathology Core (Bret M. Evers and John M. Shelton), and Transgenic Core (Robert E. Hammer)

